# Quantifying Cross-Modal Association Confidence for Single-Cell RNA-ATAC Integration

**DOI:** 10.64898/2026.05.07.723400

**Authors:** Tomoya Furutani, Hongkai Ji

## Abstract

While multimodal sequencing technologies are rapidly advancing, most single-cell and spatial datasets still measure only a single modality. Integrative computational methods for separately profiled single-cell RNA-seq (scRNA-seq) and ATAC-seq (scATAC-seq) data typically rely on the assumption that gene expression correlates with the chromatin accessibility of nearby regulatory regions. However, the strength and reliability of these correlations vary substantially across genes, and incorporating low-confidence associations can compromise integration accuracy. Here, we introduce the **CLIC** (**C**ross-modality **Li**nk **C**onfidence) score, a quantitative measure of the empirical concordance between gene expression and nearby chromatin accessibility, derived from diverse single-cell multiome datasets from the ENCODE project. CLIC scores provide prior confidence estimates for gene–peak associations across modalities. Building on this, we propose a hybrid feature selection strategy that intersects highly variable genes with high-CLIC genes, generating feature sets that better align with the assumptions of cross-modal integration methods. Across diverse publicly available single-cell and spatial datasets, and multiple state-of-the-art integration frameworks, our approach consistently improves the integration of gene expression and chromatin accessibility data, enhancing both robustness and biological interpretability.

**Graphical Abstract:** 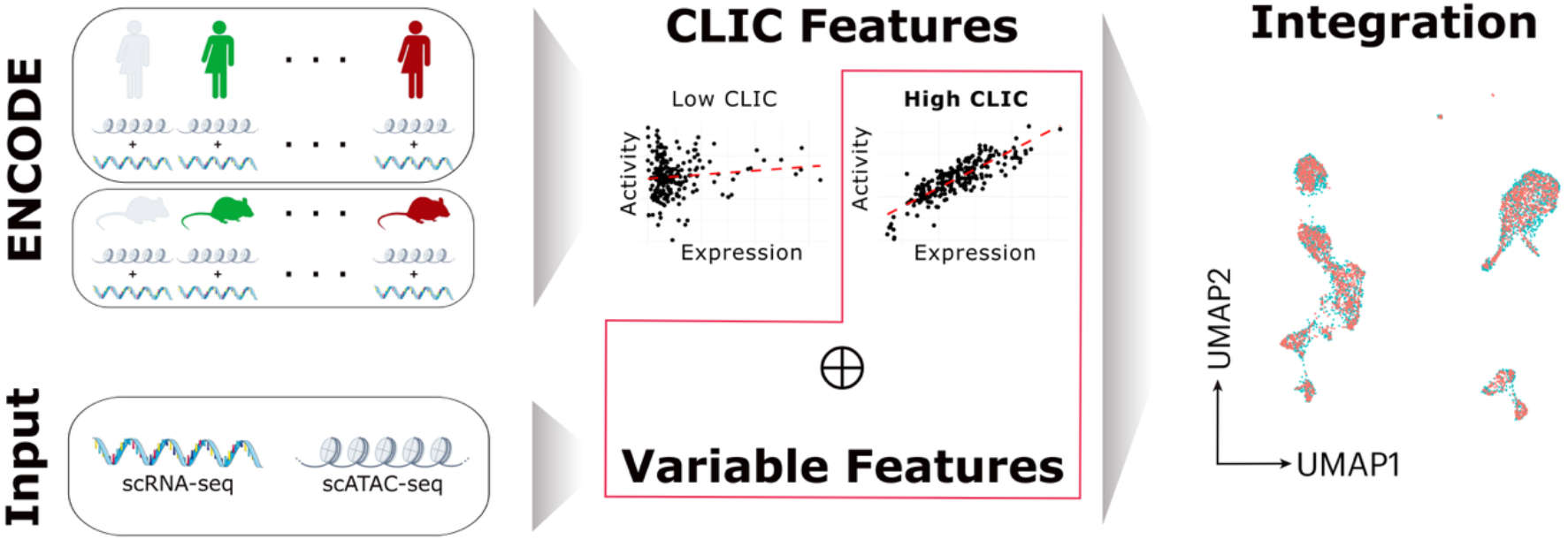

## Introduction

The emergence of single-cell omics technologies has revolutionized biological research[1, 2]. Compared to traditional bulk sequencing methods, which provide only average measurements across a cell population, measurements at a single cell resolution enable the identification of distinct cell types and states within a cell population, allowing researchers to discover complex cellular landscapes.

Biological systems are often driven by the interplay between different omics layers, such as the genome, epigenome, transcriptome, and proteome. As our understanding of complex biological phenomena grows, there is an increasing demand for a multimodal view of cellular states to identify causal relationships between different layers. In particular, single-cell RNA-seq (scRNA-seq) and single-cell ATAC-seq (scATAC-seq) which profile gene expression and chromatin accessibility, respectively, provide complementary information that, together, sheds light into cell-type specific gene-regulatory networks[3].

While assays that simultaneously profile gene expression and chromatin accessibility have emerged, these technologies often come with a variety of drawbacks[4–6]. These methods can be more complex and expensive and may yield lower data quality than single-modality measurements[7]. Thus, single-modality experiments remain widely used, and there exists a vast number of single-modality datasets, which may be leveraged to generate a multi-modal view of biological systems through cross-modality data integration. Such integration is also valuable for improving the analysis of individual data modalities. For instance, extreme sparsity of scATAC-seq data can create difficulty in cell-type annotation, whereas numerous scRNA-seq data have been well-annotated, motivating the transfer of cell-type labels from scRNA-seq to scATAC-seq data[8, 9]. Crucial to these tasks is the ability to produce high-quality integration of individually profiled scRNA-seq and scATAC-seq data.

To address this issue, numerous computational methods have been developed to integrate single-modality data[10–13]. These methods project data from separate experiments into a shared latent space, which can be used for downstream analysis such as transferring cell-type labels between modalities or generating an *in silico* multiome profile.

One primary challenge in integrating data from different modalities lies in the distinct features of each modality (genes in scRNA-seq data and chromatin-accessible regions in scATAC-seq)[14]. Many integration methods circumvent this issue by linking distinct features from the two modalities together based on prior biological knowledge. In the context of scRNA-seq and scATAC-seq integration, algorithms most commonly link the two data modalities by utilizing the regulatory relationship between genes’ nearby regulatory regions (e.g., promoters) and their transcriptional activities, assuming that gene expression and its nearby region’s chromatin accessibility is correlated. For example, Seurat[10] and LIGER[11] convert scATAC-seq data into a gene activity score matrix by summing chromatin accessibility counts in the gene body and promoter region, thereby creating matching features between scATAC-seq and scRNA-seq. Considering the gene activity score matrix as pseudo gene expression data, the two datasets are integrated via batch correction algorithms designed for scRNA-seq datasets. However, since gene activity scores do not always fully capture the complex interplay between transcription and proximal and distal regulatory elements, these scores may not correlate well with gene expression in certain genes. Including these low-correlation features can degrade the integration quality, leading to less reliable downstream analysis. Algorithms such as BindSC[12] and GLUE[13] integrate the two modalities by iteratively improving this assumption, but the initialization of feature links still relies on matching genes with their proximal peaks. Therefore, we reasoned that filtering out low-quality cross-modality feature matchings may improve the integration of scRNA-seq and scATAC-seq.

Feature selection is a fundamental preprocessing step in single-cell omics data analysis that affects the accuracy of downstream tasks[15, 16]. Current integration algorithms typically rely solely on highly variable genes as the threshold of biological relevance for feature selection. In this study, we propose incorporating an additional metric into the feature selection process based on confidence scores for feature links, computed as the empirical correlation between genes’ expression levels and local chromatin accessibility. Leveraging the extensive collection of multimodal datasets from the Encyclopedia of DNA Elements (ENCODE) consortium, we compute the CLIC (cross-modal link confidence) score as a metric to identify genes that exhibit strong linear relationships with the accessibility of their proximal peaks across biological contexts. Thus, genes with high CLIC scores can serve as a high-confidence feature link in integration with chromatin accessibility data. We select features by taking the intersection of top variable genes and top CLIC genes, thus generating inputs that are both biologically meaningful and better fit the algorithm assumptions (Figure 1a). We conduct a systematic benchmark in which we incorporate the additional preprocessing step into current state-of-the-art integration algorithms and compare their performance with that of algorithms that filter genes solely by expression variability. Across diverse datasets, scenarios, and metrics, we observed that employing the hybrid feature selection strategy improves integration quality. To facilitate easy integration into users’ existing pipelines, we have developed an R package called CLIC that implements the hybrid feature selection approach in the context of integrating scRNA-seq and scATAC-seq data.

**Fig. 1.**
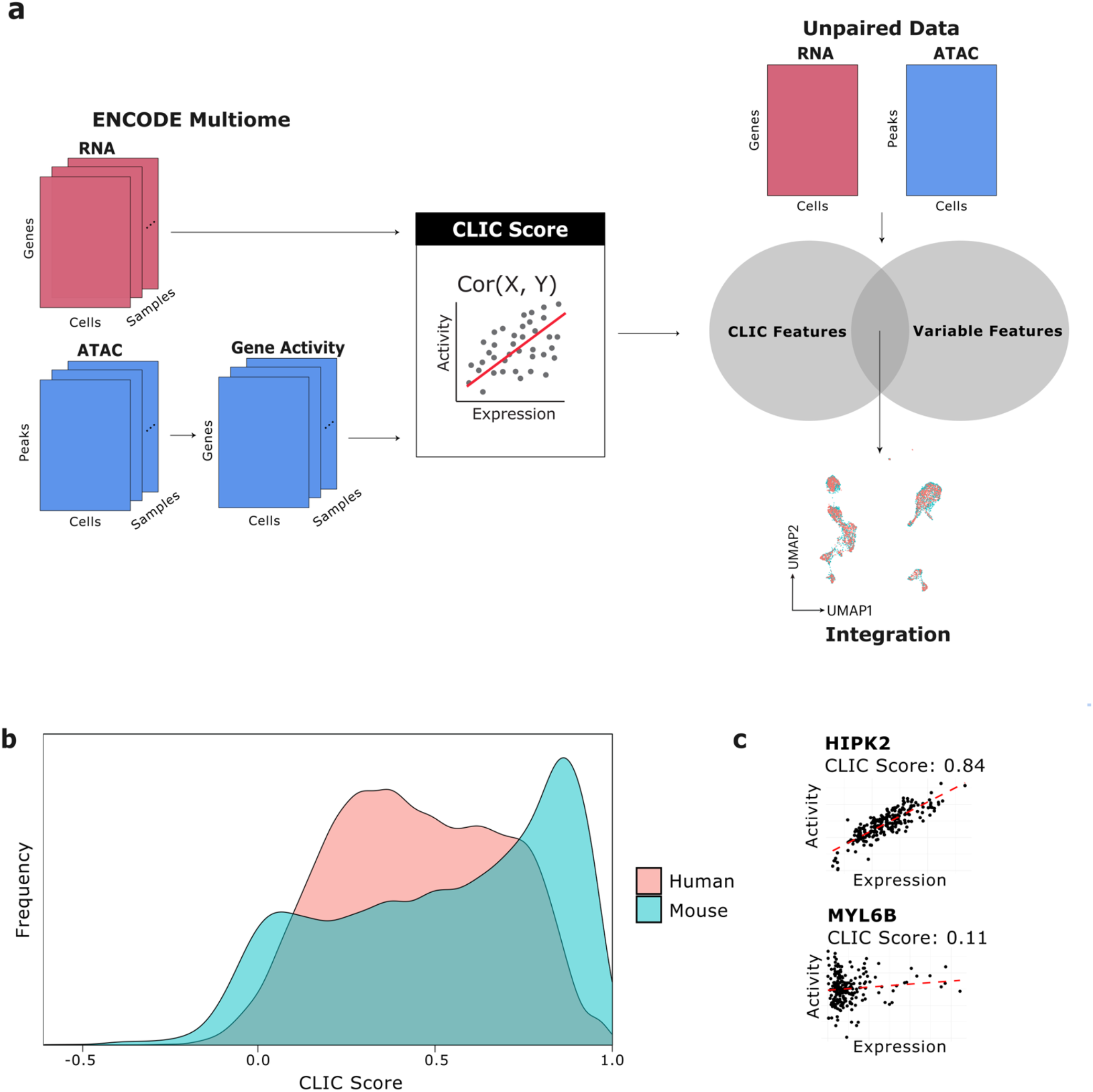
Overview of the feature selection strategy. (a) Feature selection via CLIC scores. CLIC scores are computed via the following steps. **Step 1**: Obtain single-cell scRNA-seq + scATAC-seq data from ENCODE. **Step 2**: Convert scATAC-seq data into gene activity scores and normalize them. **Step 3**: Compute Pearson’s correlation coefficient between gene expression and gene activity score for each gene. We select the intersection between the top variable genes from the gene expression modality and the highest CLIC features for integration. (b) The distribution of CLIC scores for both human and mouse. The quality of correlation between gene expression and activity varies substantially between genes. (c) Examples of high confidence feature (*HIPK2*) versus low confidence feature (*MYL6B*). Each data point represents a pseudobulk sample.

## Materials and Methods

### Hybrid Feature Selection Using CLIC Scores

#### Preprocessing ENCODE Multiome Data

To quantify the correlation between expression level and local chromatin accessibility for each gene, we leverage an atlas of single-cell multiome datasets that co-profile gene expression and chromatin accessibility, obtained from ENCODE. For human, we obtained 48 samples from 13 tissues, totaling 486,455 cells. For mouse, we obtained 40 samples from 5 tissues, totaling 333,019 cells. We downloaded the raw sequencing files from the ENCODE portal (encodeproject.org)[17] and processed them using Cell Ranger ARC v2.0.1[18]. The list of all samples used in this study and their sequencing metrics can be found in the supplementary material, and on our GitHub repository (https://github.com/oldvalley49/CLIC_compute). The gene expression counts were processed using the standard Seurat pipeline. Gene activity score was computed for each cell by counting fragments overlapping the gene body and a 2-kb upstream region of each gene using the GeneActivity() function in Signac. The cells were filtered using the following criteria: 500 < nCount_ATAC < 60,000, 500 < nCount_RNA < 40,000, nucleosome_signal < 2, TSS.enrichment > 1, and percent.mt < 10.

#### Computation of CLIC Scores

For human and mouse data, respectively, samples are integrated within their tissue types by Harmony[19] based on gene expression data. The integrated objects are individually clustered using the Louvain algorithm with a resolution of 0.5. We aggregated the gene expression counts and gene activity scores for each cluster, creating bimodal, paired pseudobulk data. The pseudobulked data represent the interaction between expression and activity score for each gene across various tissues and cell types. We used 233 and 144 pseudobulk samples for human and mouse, respectively, to compute CLIC scores.

For both modalities, library size normalization is performed, followed by a ln(*x*+1) transformation and quantile normalization. For each gene, CLIC scores are computed as Pearson’s correlation coefficient between its normalized gene expression and gene activity score.

#### Hybrid Feature Selection by Combining Gene Variability and CLIC Score

To select *n* genes, we first define *m* as a hyperparameter, where *m* is strictly larger than *n*. We identify the top *m* most variable genes using the FindVariableFeatures() function in Seurat. Next, we select the top *p* genes based on their CLIC score, calibrating *p* such that the intersection of the *m* most variable features and top *p* genes with the highest CLIC scores yields exactly *n* genes. This final set of *n* genes is used for integration.

## Benchmarking Pipeline

### Benchmarked Integration Algorithms

To compare our CLIC-based feature selection algorithm to baseline variable feature selection, we incorporated the two different methods into the workflows of four well-known algorithms for multimodal integration, Seurat v3[10], LIGER[11], bindSC[12], and GLUE[13]. Seurat v3, LIGER, and bindSC were run using R packages ‘Seurat’ (v.5.2.0), ‘rliger’ (v.2.1.0), ‘bindSC’ (v.1.0.0). GLUE was ran using the Python package ‘scglue’ (v.0.3.2). We ran each method according to the recommended preprocessing steps and hyperparameters. A total of 2000 genes were selected as features for integration across all datasets and scenarios. For Seurat, LIGER, and bindSC, the gene activity matrix was computed by summing fragments overlapping each gene body and 2kb upstream of the transcription start site. For GLUE, a prior regulatory graph is constructed instead of a gene activity matrix, with genes and peaks as nodes and edges between gene-peak node pairs within 2kb of each other. Therefore, a node representing an ATAC peak is connected to a gene node if it overlaps with the gene body or 2kb upstream of the transcription start site. For datasets in which fragment files were not available, gene activity scores were computed by summing signals from peaks overlapping each gene body and promoter region.

#### Seurat v3

Seurat[10] takes two inputs: the gene expression count matrix, as well as the gene activity matrix, computed by summing fragments from scATAC-seq overlapping each gene and its promoter region. Seurat then identifies a set of anchors between the gene expression and gene activity matrix through canonical correlation analysis (CCA), which is then used to transfer either discrete labels or continuous data from one modality to the other.

#### LIGER

LIGER[11] uses integrative non-negative matrix factorization (iNMF) to embed unpaired multi-omics datasets into a joint latent space. As iNMF relies on shared features across the unpaired dataset, scATAC-seq data is converted to a gene activity matrix.

#### BindSC

BindSC[12] uses the gene expression counts matrix, peak-by-cell matrix, and gene activity matrix as input. Gene activity matrix is used to initialize the “modality fusion matrix”, which links the distinct features of the two modalities, which is then iteratively improved via bi-order canonical correlation analysis CCA.

#### GLUE

GLUE[13] aligns cells across the two modalities via a guidance graph, where each vertex represents a feature and each edge implies a regulatory relationship. The guidance graph is initialized using prior biological knowledge; for scRNA-seq and scATAC-seq integration, ATAC peaks are connected to genes with a positive edge if they overlap in either the gene body or promoter region.

### Benchmark Datasets

To benchmark our method’s performance, we gathered publicly available single-cell and spatial datasets in which gene expression and chromatin accessibility were co-profiled. A full list of the data used in this study is provided in Supplementary Table 1. All datasets were converted into common formats, including an RNA gene-by-cell count matrix, an ATAC peak-by-cell count matrix, a gene activity matrix, and the cell-type labels.

Notably, the cell-type annotation for the 10X PBMC data was obtained by following the Seurat tutorial[20]. Similar cell types were then joined together to create 9 high-level cell-type labels to be used for benchmarking, as done in a recent benchmarking study: B-cells (“B”), CD4 T cells (“CD4 T”), CD8 Naïve T cells (“CD8 Naïve”), CD8 Effector Memory T cells (“CD8 TEM”), Dendritic cells (“DC”), Monocytes (“Mono”), Nature killer cell (“NK”), other T cell (“other_T”), and other cell categories (“other”)[21].

For all other datasets, cell-type labels were published by the original authors.

### Evaluation Criteria

One way to evaluate method performance is to use multiome data with paired scRNA-seq and scATAC-seq from the same cell while pretending that the two modalities were separately profiled from different cells. In such a case where ground-truth pairs are known, an accurate integration should map them close to each other in the latent space. Additionally, the generated latent space should preserve proximity within each cell type while mixing cells from the two modalities in a homogeneous manner. Accordingly, we evaluated the integration performance using the following criteria

**FOSCTTM (Fraction of Samples Closer than True Match)** was used to measure single-cell-level alignment accuracy[22]. FOSCTTM requires the knowledge of ground-truth cell-to-cell correspondence. Thus, it was computed when benchmarking using paired data.

Given two datasets, both of size *N*, such that the *i*th cells in the two datasets are paired together, we compute FOSCTTM as follows:

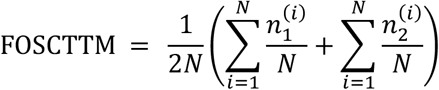

where 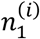 and 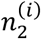 are the number of cells in the first and second datasets, respectively, that are closer to the *i*th cell than their true matches in the other dataset, determined using Euclidean distance in the latent space. FOSCTTM is then transformed as 1 − *FOSCTTM* so that higher values indicate better performance. FOSCTTM ranges from 0 to 1.

**ASW (Average Silhouette Width)** was used to measure cell-type separation in the latent space. ASW requires ground-truth cell-type labels for each dataset[23]. For each cell, we compute the silhouette score:

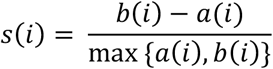

where *a*(*i*) is the average intra-cell-type distance and *b*(*i*) is the minimum average distance to cells in any other cell type. The average silhouette width over all cells is computed as follows:

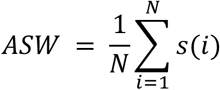

The computed scores are then transformed using (x+1)/2 to set the ASW range to 0 to 1. Higher values indicate better cell-type separation.

**Batch ASW (Batch Average Silhouette Width)** measures how well cells from different batches mix within each cell type[23, 24]. It is based on the average silhouette width (ASW) computed using batch labels, where 0 indicates perfect mixing. To make higher scores reflect better mixing, the ASW is subtracted from 1. Batch ASW is first calculated per cell type as follows:

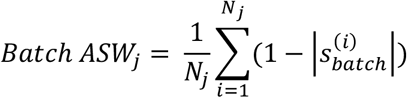

where 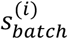 is the silhouette width for the *i*th cell computed using batch labels, and *N*_*j*_ is the number of cells in cell type *j*. Then, they are averaged across all cell types to get the final score:

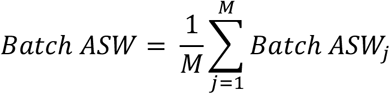

where *M* denotes the number of unique cell type labels. A score of 1 indicates perfect mixing; 0 indicates strong batch separation.

**ARI (Adjusted Rand Index)** was used to evaluate label-transfer accuracy. For each method, the default function for cell-type label transfer is used if such a function is implemented. If not, the k-nearest neighbors (k-NN) algorithm with k=30 is used to transfer labels from the RNA modality to the ATAC modality. Higher ARI means better accuracy in cell-type label transfer.

**kNN-AUC (k-Nearest Neighbors Area Under the Curve)** was used to assess how well the local neighborhoods of cells align between scRNA-seq and scATAC-seq modalities after integration[25]. Let *f*(*i*) ∈ *F* be the scRNA-seq measurement of cell *i*, and *g*(*i*) ∈ *G* be the scATAC-seq measurement of cell *i*. kNN-AUC is computed as the average percentage overlap of neighborhoods of *f*(*i*) in *F* with neighborhoods of *g*(*i*) in *G*, with higher values indicating better local structure preservation. In our analysis, we define neighborhoods using the top 1% of closest neighbors.

## Results

### Framework for Hybrid Feature Selection Using CLIC Scores

Based on available data, we computed CLIC scores for human and mouse genes. As expected, the level of correlation between gene expression and local chromatin accessibility varied substantially across genes, showing that treating all genes equally in integrating the two modalities may introduce bias (Human: Min = -0.31, Mean = 0.43, Max = 0.96; Mouse: Min = -0.62, Mean = 0.50, Max = 0.99) (Figure 1b).

Given a total of *n* features to select for integration and a hyperparameter *m* (with *m* > *n*), we calibrate *p* such that the intersection between the top *m* most variable features and the top *p* features with the highest CLIC score yields exactly *n* features. These selected features are then used for the downstream integration task. The hyperparameter *m* controls the relative weighting between expression variability and correlation in feature selection, with larger values of *m* placing greater emphasis on the CLIC score.

### Comprehensive Benchmark Using Paired scRNA-seq and scATAC-seq Data

To evaluate the performance of the hybrid feature selection strategy in cross-modal integration, we first incorporate CLIC into the workflows of four current state-of-the-art integration methods, which use proximity between peaks and genes as prior knowledge for cross-modal alignment[10–13]. We then compiled datasets generated from simultaneous profiling of gene expression and chromatin accessibility within the same cell along with cell-type annotations. These datasets comprise three human tissues and three mouse tissues in blood, bone marrow, brain, skin, and retina (Supplementary Table 1). The bone marrow mononuclear cells (BMMC) data consist of multiple batches from four different sites and nine different donors. Here, to avoid complexity caused by intra-modality batch effects, we select two batches with the greatest number of cells that share neither the site nor the donor (s1d2 and s4d8). We treat the data from the two modalities as originating from two separate single-modality experiments and evaluate the integration quality using the ground-truth pairing information.

We used the following metrics to evaluate integration performance:

- **FOSCTTM (Fraction of Samples Closer Than the True Match)**: Measures how close the ground-truth pairs are to each other relative to other cells[22].
- **ASW (Average Silhouette Width)**: Measuring the separation of different cell types using ground-truth cell type annotation[23].
- **Batch ASW (Batch Average Silhouette Width)**: Measuring how well the two modalities are mixed in the latent space[23].
- **ARI (Adjusted Rand Index)**: Measures cell-type prediction accuracy when cell-type labels are transferred from scRNA-seq data to scATAC-seq data, accounting for randomness.
- **kNN-AUC (k-Nearest Neighbors Area Under the Curve)**: Measures how well the local neighborhoods of cells align between scRNA-seq and scATAC-seq modalities after integration. A higher score indicates better preservation of local cell neighborhoods[25].

To facilitate interpretation, FOSCTTM was transformed so that higher values indicate better performance for all metrics used in the benchmark. Specifically, since FOSCTTM originally ranges from 0 to 1, with lower values indicating better performance, we compute the transformed FOSCTTM as 1-FOSCTTM. Cell type annotations used for benchmarking were published by the original authors for all datasets, except PBMC. PBMC data were annotated following Seurat’s tutorial on weighted nearest neighbor analysis[20], using both RNA and ATAC modalities for annotation, and processed further to join similar cell types, as done in a recent benchmarking study[21]. For a fair method comparison, a total of 2000 genes were selected for the variable feature selection and the hybrid feature selection strategy. We also vary the weight given to the CLIC score in hybrid feature selection to assess the method’s robustness to hyperparameter settings.

Each algorithm-dataset pair is tested using five feature selection methods, including the default variable feature selection (baseline) and a hybrid strategy with four different values of hyperparameter *m*, denoted as CLIC_m (CLIC_4000, CLIC_5000, CLIC_6000, and CLIC_8000). Each pair of integration algorithm and metric is used as a criterion to evaluate the feature selection method. For each criterion, methods were ranked according to the average score across the benchmark datasets. Finally, we ranked each method according to its average rankings over all criteria. This amounts to evaluating a total of 700 different combinations (5 feature selection methods x 7 datasets x 4 integration algorithms x 5 metrics).

Overall, integration algorithms performed better when the input features were selected using the hybrid strategy (Figure 2). Across integration algorithms and metrics, filtering cells based on their correlation robustly improved integration quality in most cases and parameter settings.

**Fig. 2.**
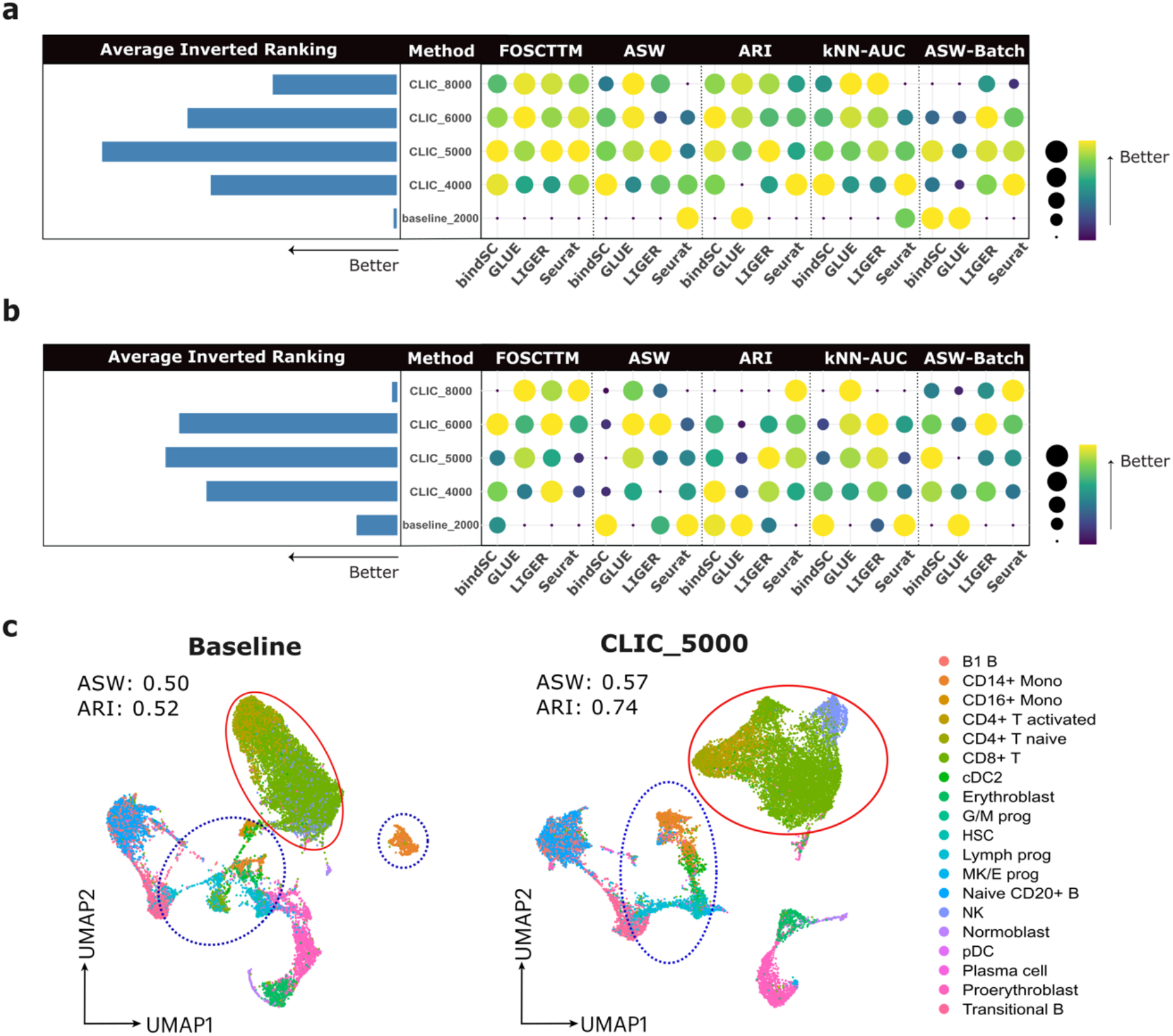
Benchmarking result on paired scRNA-seq + scATAC-seq data (a) Performance comparison between baseline variable feature selection and hybrid feature selection on human dataset. Each column represents an algorithm metric combination, with the results averaged across all human datasets. The bar plot represents the average inverted ranking of all feature selection strategies. (c) Performance comparison, with the results averaged across all mouse datasets. (c) UMAP visualization of the latent space generated by LIGER from BMMC-s4d8 data.

By comparing performance across individual datasets, we found that the hybrid methods outperformed the baseline feature selection approach in most datasets, highlighting their ability to enhance cross-modal integration both effectively and efficiently (Figure S1). The performance gains were especially pronounced in more complex datasets, such as BMMC and TDBM, compared to PBMC, suggesting the hybrid approach’s potential in handling challenging integration scenarios. In minority cases where the baseline method outperformed the hybrid approach, the hybrid method performed comparably to the baseline method, with the decrease in performance being much less significant than the increase in performance in other cases. As a representative example, we examined the resulting joint UMAP embedding from running LIGER on the BMMC dataset (Figure 2c). When features were selected via CLIC, LIGER produced a clearer separation of different cell type clusters. These improvements in visual separation aligned with the quantitative gains observed in our cell-type separation metrics (ASW, ARI), providing visual confirmation that the CLIC features enhance LIGER’s ability to recover biologically meaningful structure in complex single-cell datasets.

### CLIC Improves Integration of Unpaired scRNA-seq and scATAC-seq Data

While ground-truth pairing information from multiome datasets is valuable for benchmarking, it reflects an idealized scenario without batch effects between modalities. In realistic cross-modal integration settings, gene expression and chromatin accessibility data are often generated from different donors and laboratories, introducing substantial batch effects. To evaluate algorithms under more practical conditions, we leverage the internal batch structure of the BMMC dataset. Specifically, we selected three batch pairs that differ in both donor and originating lab and performed computational integration of the data (Figure 3a). These pairs span cases where the two modalities contain either comparable or imbalanced numbers of cells. Since the ground-truth pairing information is no longer available, we assessed integration performance using only cell type-level metrics (ASW, Batch ASW, and ARI). This amounts to evaluating a total of 180 different combinations (5 feature selection methods x 3 dataset pairs x 4 integration algorithms x 3 metrics).

**Fig. 3.**
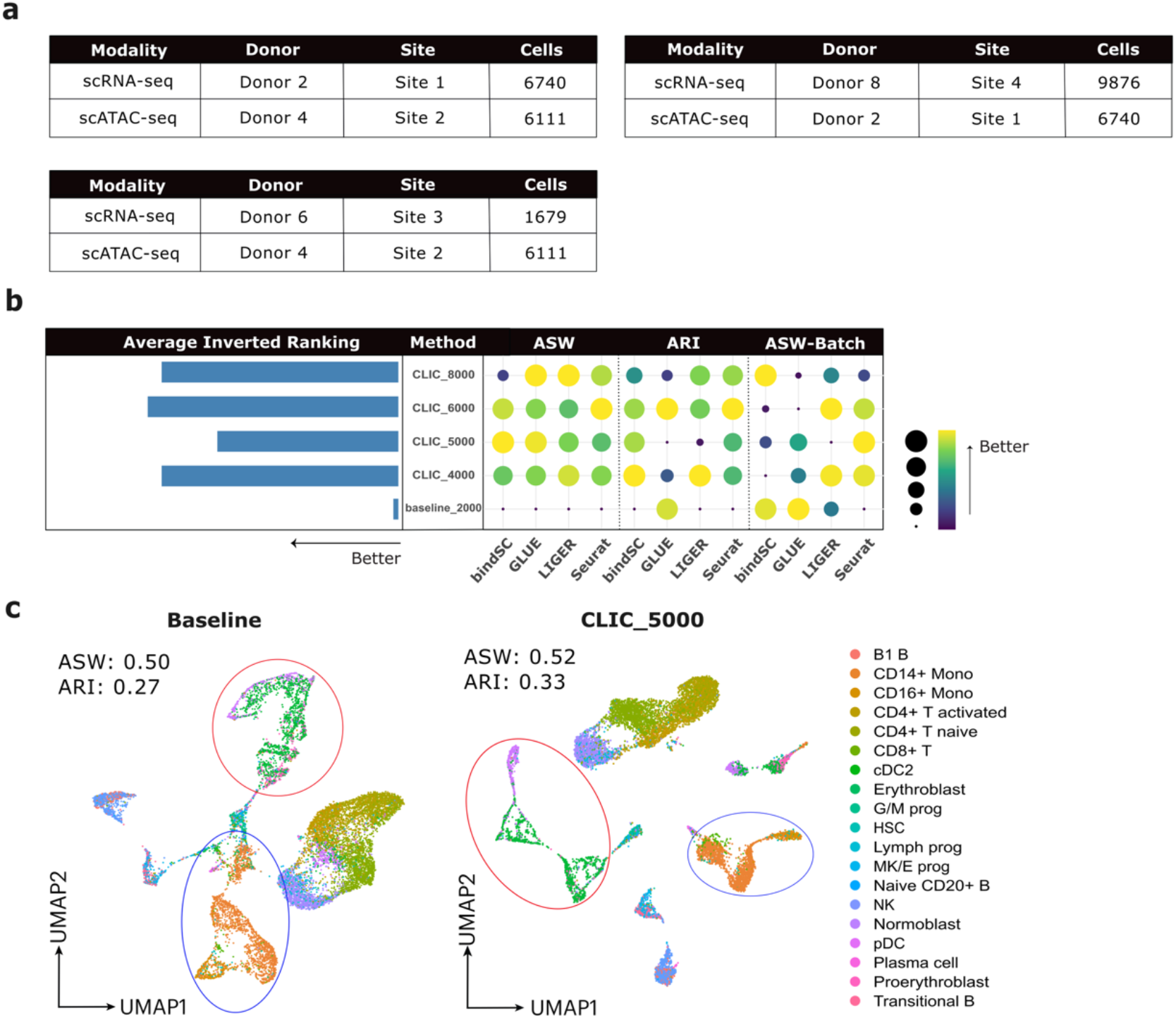
Benchmarking result on unpaired BMMC data. (a) Batches used in the benchmarking experiment. The two batches do not share a donor or an experiment site. (b) Performance comparison between baseline variable feature selection and hybrid feature selection on the unpaired BMMC dataset. Each column represents a combination of algorithm metrics, with the results averaged across all human datasets. The bar plot represents the average inverted ranking of all feature selection strategies. (c) UMAP visualization of the latent space generated by LIGER from integrating scRNA-seq data from BMMC-s1d2 and scATAC-seq data from BMMC-s2d4.

Again, incorporating the CLIC score into feature selection led to robust improvements in integration quality (Figure 3b, Figure S2). Notably, we observed enhanced cell type separation and improved label transfer, highlighting the potential of CLIC to improve downstream analyses of single-cell data. Visualizing the generated UMAP embedding from LIGER, we observed a clearer organization of cell types, accompanied by increases in numerical metrics (Figure 3c).

### CLIC Improves Integration of Spatial Transcriptomics and Chromatin Accessibility Data

Recent advancements in spatial technologies have revolutionized our ability to investigate cellular processes while preserving spatial context. Spatial transcriptomics, which enables measurement of gene expression with spatial resolution, has already been commercialized and is becoming increasingly prevalent in research^23^.

The integration between spatial omics data and single-cell omics data offers the potential to provide a comprehensive view of gene regulation in a spatial context. This integration could reveal how chromatin accessibility patterns correlate with gene expression across different tissue regions, potentially uncovering spatially dependent regulatory mechanisms.

Recently, datasets from spatial co-profiling of gene expression and chromatin accessibility have become available[28–30]. However, the technology for simultaneous spatial profiling of both modalities is not yet readily accessible. As such, cross-modal integration algorithms designed for single-cell datasets are useful for leveraging more widely available single-modality spatial dataset to gain insights into gene regulatory patterns across tissue spaces.

To test our method for improving cross-modal integration in a spatial context, we leverage the recently published data from spatial co-profiling of gene expression and chromatin accessibility[28–30]. Specifically, we tested five spatial RNA–ATAC datasets from human (brain and skin) and mouse (brain and two embryonic samples), generated using three distinct technologies (spatial ATAC-RNA-seq, spatial-Mux-seq, and Slide-tags). We then implement an equivalent benchmarking workflow to that of paired single-cell data, in which we consider the data as originating from two different single-modality experiments—spatial RNA-seq and single-cell ATAC-seq—and use their ground-truth cell-to-cell correspondence for calculating the integration metric (FOSCTTM, kNN-AUC). We did not use cell type-level metrics (ASW, Batch ASW, and ARI) as they require ground-truth cell type annotation, which was not available. This amounts to evaluating a total of 200 different combinations (5 feature selection methods x 5 datasets x 4 integration algorithms x 2 metrics).

In four of five spatial datasets, feature selection using CLIC scores outperformed the baseline method in both metrics, highlighting its ability to improve cross-modal integration in a spatial context (Figure 4a, 4b, S3). One application of multimodal integration algorithms is gene regulatory network inference, achieved by aligning spatial gene expression and single-cell chromatin accessibility. Seurat supports this by generating a pseudo–gene expression profile from chromatin accessibility data. To illustrate the potential impact on such downstream tasks, we imputed the expression level of a granule cell layer (GCL) marker gene *THY1* and visualized its spatial distribution, and saw that spatial gene expression pattern was better predicted by ATAC-seq when CLIC scores were considered (Figure 4c).

**Fig. 4.**
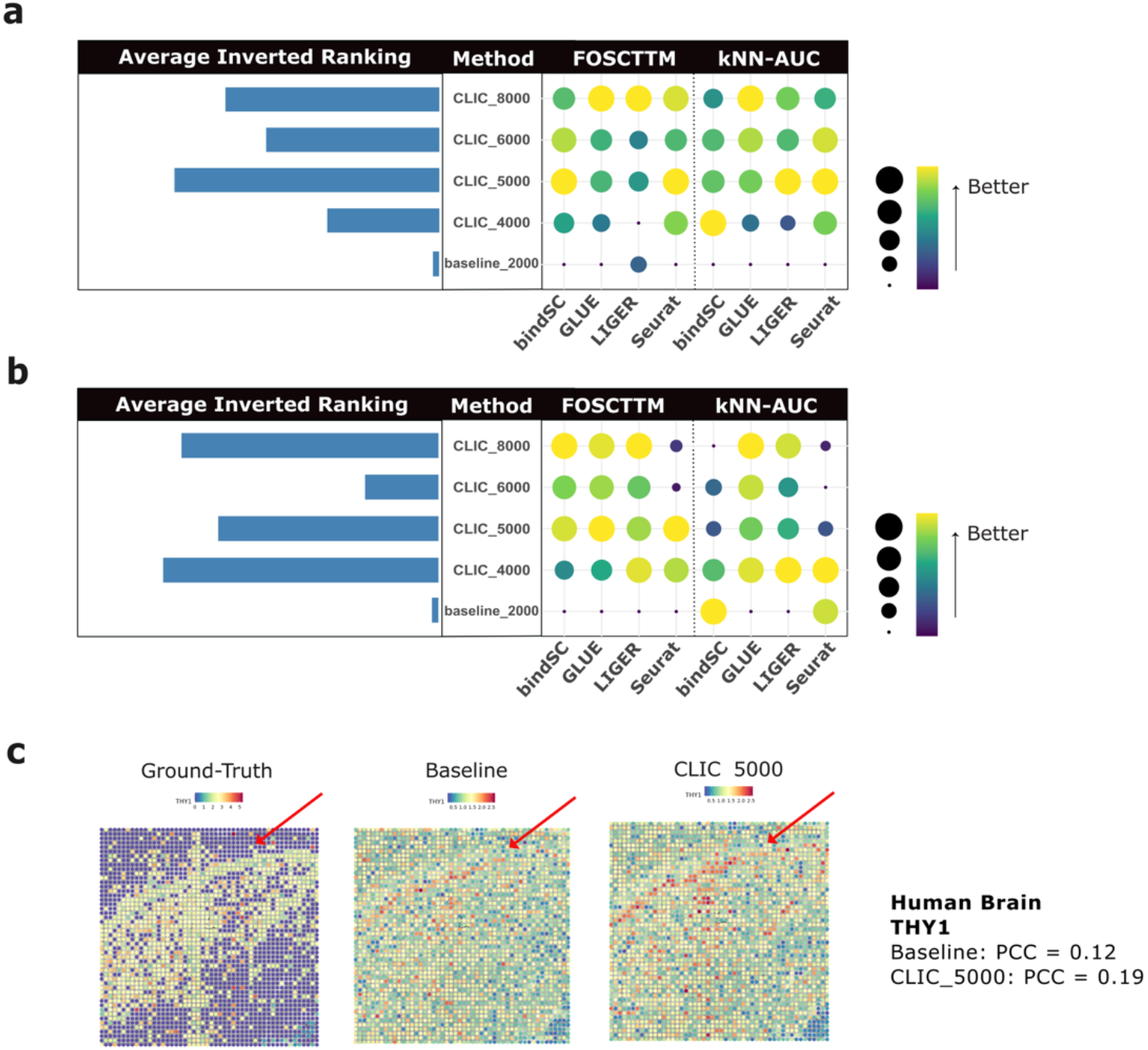
Benchmarking result on multiome spatial data. (a) Balloon plot of benchmarking results from human spatial data. (b) Balloon plot of benchmarking results from mouse spatial data. (c) Comparison of the spatial distribution of ground-truth gene expression, imputed gene expression via default Seurat settings, and imputed gene expression when filtered via human CLIC score. *THY1* is a known marker of the granule cell layer (GCL).

### CLIC Is Robust to Hyperparameter Choices and Gene Activity Models

Finally, we examined the robustness of our feature selection method to methodological choices that may vary across experimental settings. Specifically, we analyzed how changes in hyperparameter selection and differences in gene activity score computation influence the observed improvement in integration quality.

A natural question for practical use of the package is how to select the hyperparameter *m*. This parameter controls the relative weight given to the CLIC score versus expression variability, with higher values of *m* placing more emphasis on genes with higher CLIC scores. As shown in our benchmarks, the method was robust to changes in *m* within the range of 4,000–8,000. We further evaluated the algorithm with a wider range of hyperparameters on single-cell multimodal datasets and found that in the majority of cases, FOSCTTM scores improved for *m* values between approximately 2,500 and 10,000 when selecting 2000 features in total (Figure 5a). Values outside this range tended to either underweight the CLIC score or yield biologically uninformative features by sacrificing too much expression variability. For usability, in our package, which is interfaced with Seurat, we set the default *m* value to 6,000 in CLIC, the optimal empirical parameter when using Seurat. However, users may freely change this value.

**Fig. 5.**
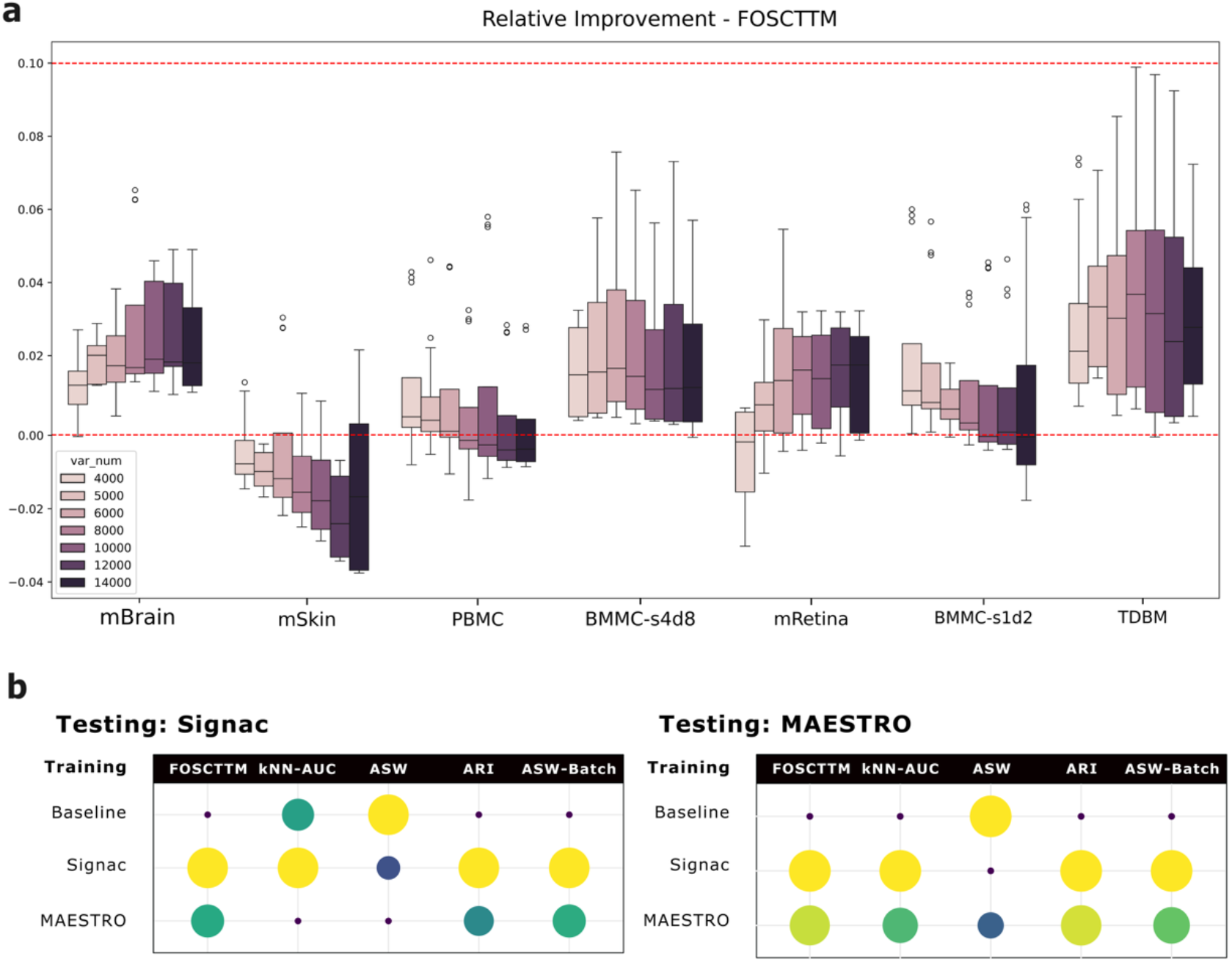
Sensitivity analysis on hyperparameter *m* and activity model selection. (a) Relative improvement in FOSCTTM score from baseline across various hyperparameter settings. Horizontal red line at 0 indicates baseline performance. (b) Balloon plot of benchmarking performance using hybrid feature selection versus baseline over all CLIC parameter choices and benchmarked integration algorithms. Testing activity models are used for input into integration algorithms, and training activity models are used to compute CLIC scores.

Another important consideration is the choice of gene activity model, which influences how chromatin accessibility is mapped to gene-level signals. In our benchmark, we primarily used Signac[26]’s approach, which quantifies gene activity by summing ATAC fragments that overlap each gene body and its promoter region, effectively capturing accessibility of nearby regulatory elements. An alternative widely used model is implemented in MAESTRO[31], which weights distal chromatin regions using an exponential decay function based on genomic distance; accessibility signals closer to a gene’s transcription start site contribute more heavily, whereas distal elements have progressively lower influence. To evaluate whether our feature selection method is robust to such modeling choices, we performed a sensitivity analysis across gene activity definitions. Gene activity computation appears in two stages of our workflow: (i) during training, where ENCODE data are used to compute CLIC scores, and (ii) during testing, where gene activity inputs are fed into multimodal integration algorithms. We therefore evaluated all four possible combinations of gene activity models across training and testing (e.g., training with MAESTRO and testing with Signac, and vice versa). We observed that improvements in integration quality were largely preserved. However, CLIC scores computed based on Signac consistently outperformed both baseline and CLIC scores computed using MAESTRO. Therefore, we have set the default CLIC score for our package to be those computed using Signac.

## Discussion

In summary, we present a hybrid feature selection strategy for the computational integration of scRNA-seq and scATAC-seq data. In contrast to the traditional method of selecting genes solely by their expression variability, we also consider the confidence level of feature links evaluated by the degree of correlation between local accessibility and expression for each gene. CLIC scores are computed using human and mouse multimodal data obtained from ENCODE. Our approach addresses the key limitation in current integration algorithms: their underlying assumption that closely located regions and genes have high correlation may not be true.

Our benchmarking results demonstrate that a hybrid feature selection strategy outperforms highly variable feature selection across diverse scenarios, integration methods, datasets, and metrics. To facilitate the integration of this method into users’ existing data analysis pipelines, we developed the R package CLIC, which implements the hybrid feature selection strategy.

Given our framework, which leverages publicly available multiome data, the computed CLIC scores can be expected to improve and can be expanded to other species as more data become available in the future. While our focus has been on feature selection in the computational integration of scRNA-seq and scATAC-seq data, we expect the framework for improving prior knowledge assumption in cross-modality integration using publicly available data to be generalizable across various modalities.

Beyond improving cross-modal integration, this study highlights the benefits of public atlas data as valuable priors for analyzing new datasets. Curating and expanding these large, diverse collections of biological conditions is essential for enabling the community to build more accurate, generalizable, and biologically grounded computational models.

## Additional Files

Supplementary Figures:

- Supplementary Figure S1: Improvement in each metric from the benchmark using paired data.
- Supplementary Figure S2: Improvement in each metric from the benchmark using unpaired data.
- Supplementary Figure S3: Improvement in each metric from the benchmark using spatial data.

Supplementary Tables:

- Supplementary Table S1: List of benchmarking datasets
- Supplementary Table S2: List of ENCODE human datasets used to compute CLIC scores and their sequencing metrics
- Supplementary Table S3: List of ENCODE mouse datasets used to compute CLIC scores and their sequencing metrics

## Acknowledgments

The authors would like Steven Liu from the Ji Lab and Ziqi Fu from Harvard University for sharing some processed datasets.

## Availability of Data and Materials

The data used in this study are all publicly available. The ENCODE data used to compute CLIC scores, along with their sequencing metrics, are available in Supplementary Tables 2 and 3. The code for preprocessing the data and computing CLIC scores is available under https://github.com/oldvalley49/CLIC_compute.

Additional information of datasets used for benchmarking (PBMC[32], BMMC[33], TDBM[34], mSkin[35], brain[28], Melanoma[29], mBrain[35], mBrain2[30], mRetina[12], mEmbryo[28], mEmbryo2[30]) are included in Supplementary Table 1. Code to reproduce the benchmark analysis in the study can be found under https://github.com/oldvalley49/CLIC_benchmark.

## Availability of Software

The R package CLIC is available on GitHub at https://github.com/oldvalley49/CLIC and on Zenodo at https://doi.org/10.5281/zenodo.18966333

## Ethics Approval and Consent to Participate

Not applicable.

## Consent for Publication

Not applicable.

## Author Contributions

Conceptualization, H.J.; Methodology, Software, Formal Analysis, Investigation, Data Curation, Visualization, Writing – Original Draft, T.F.; Resources, Supervision, Funding Acquisition, H.J.; Writing – Review & Editing, T.F. and H.J.

---

**Fig. S1.**
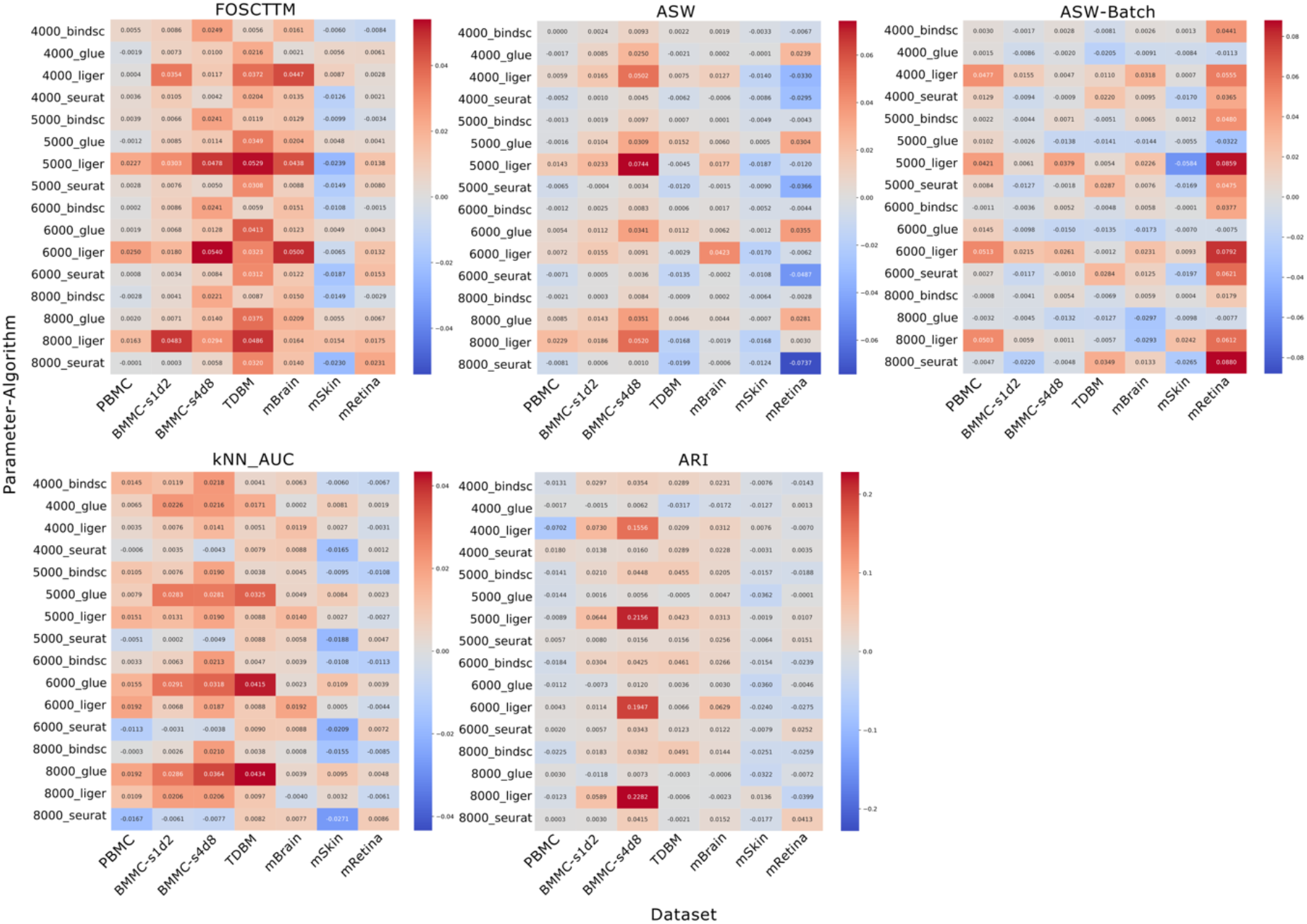
Improvement in each metric from benchmarking experiment using paired data. Each row of the heatmap represents a combination of the parameter *m* for CLIC_m and the integration algorithm used. Each column represents the dataset. Red represents improvement over baseline.

**Fig. S2.**
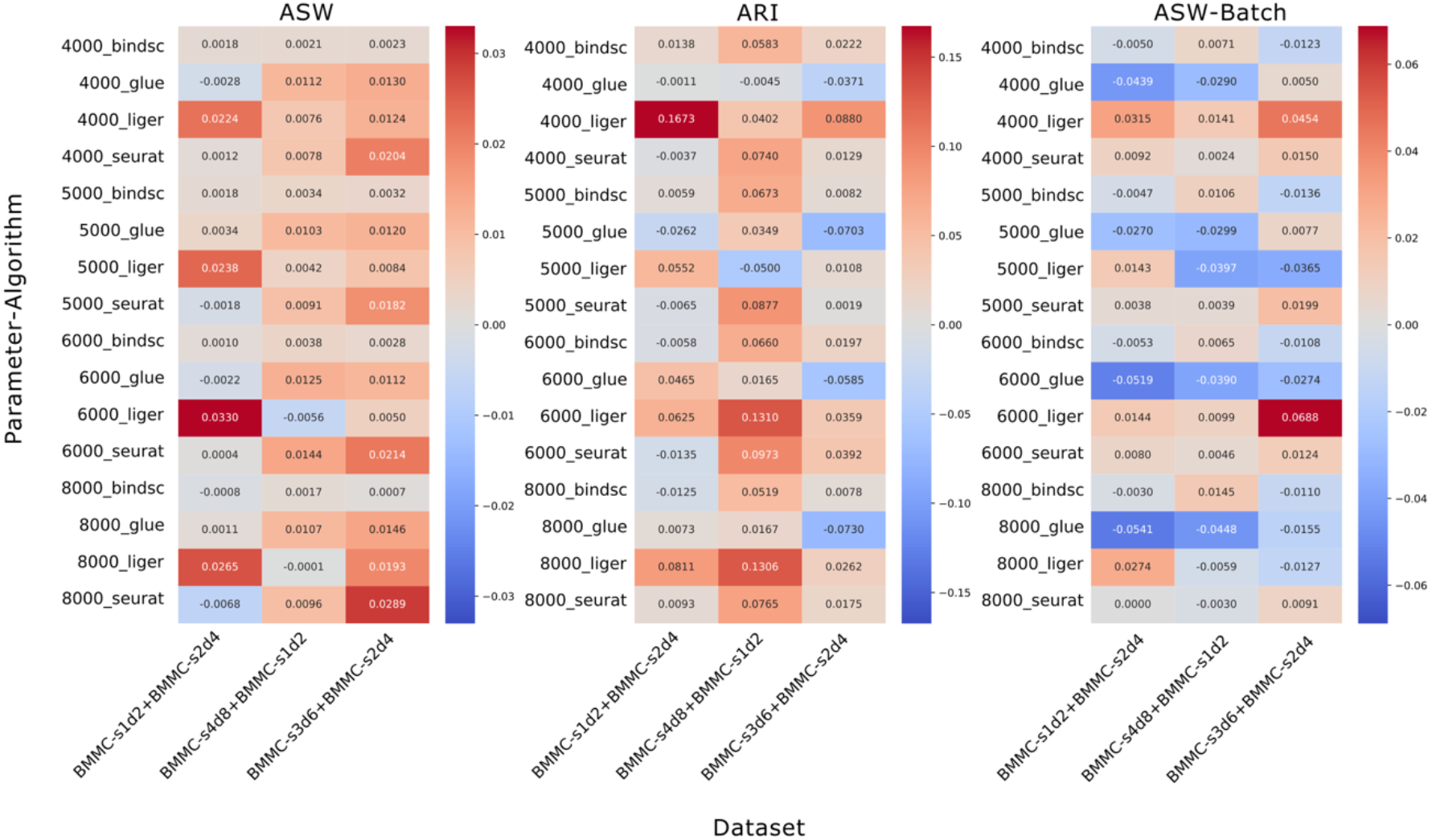
Improvement in each metric from benchmarking experiment using unpaired BMMC data. Each row of the heatmap represents a combination of the parameter *m* for CLIC_m and the integration algorithm used. Each column represents the pair of batches used (RNA+ATAC). Red represents improvement over baseline.

**Fig. S3.**
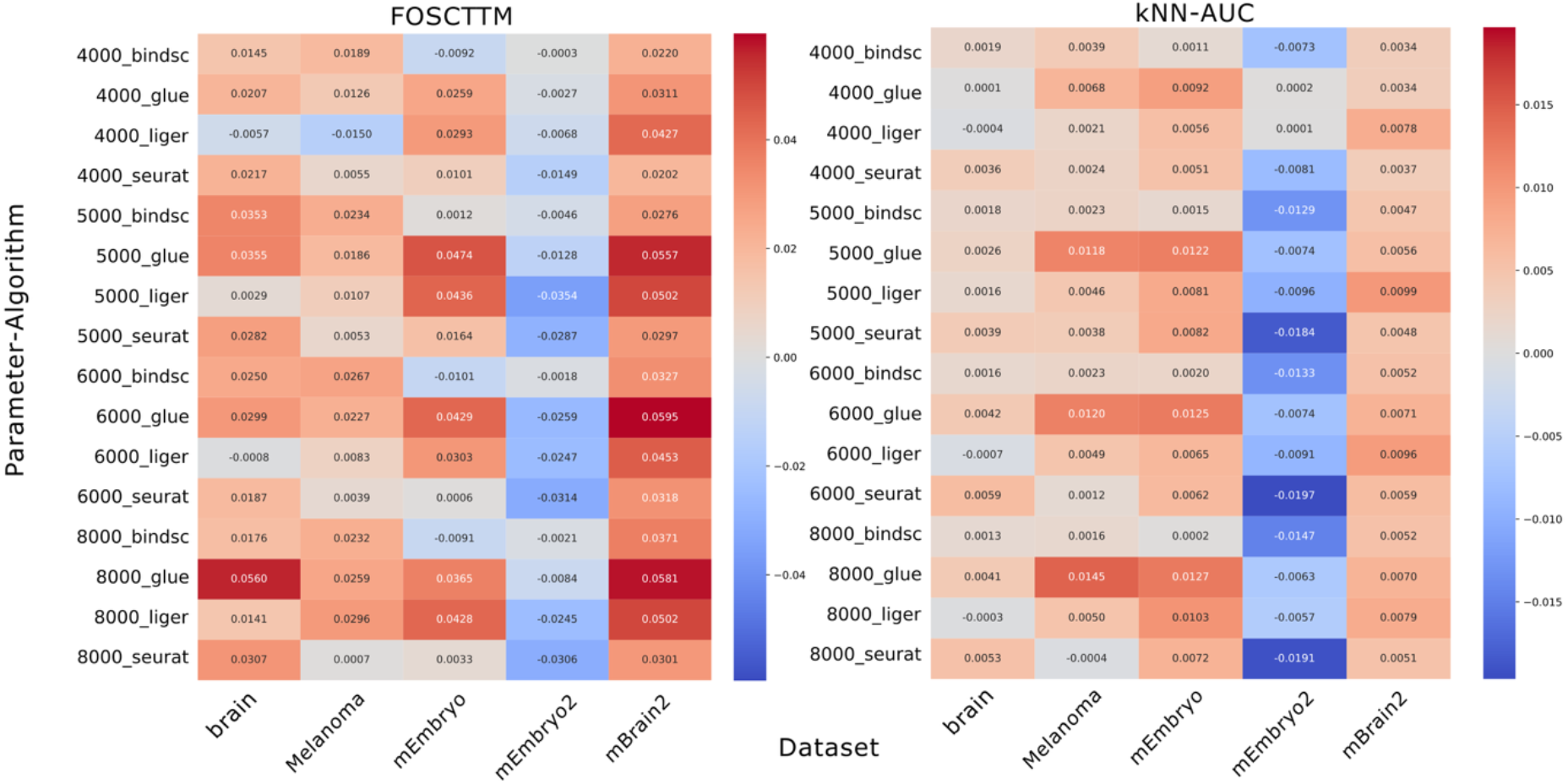
Improvement in each metric from benchmarking experiment using spatial multiome data. Each row of the heatmap represents a combination of the parameter *m* for CLIC_m and the integration algorithm used. Each column represents the spatial dataset. Red represents improvement over baseline.

